# Genetic Algorithms as a method to study adaptive walks in biological landscapes

**DOI:** 10.1101/2020.07.29.226324

**Authors:** Edith Invernizzi, Graeme D Ruxton

**Affiliations:** Sir Harold Mitchell Building, School of Biology, University of St Andrews, KY16 9TH, St Andrews, United Kingdom; Dyers Brae Building, School of Biology, University of St Andrews, KY16 9TH, St Andrews, United Kingdom

**Keywords:** evolutionary modelling, evolutionary theory, adaptation, evolutionary algorithm, Sir Philip Sidney game

## Abstract

The metaphor of fitness landscapes is common in evolutionary biology, as a way to visualise the change in allele or phenotypic frequencies of a population under selection. Understanding how different factors in the evolutionary process affect the trajectory of the population across the landscape is of interest to both theoretical and empirical evolutionary biologists. However, fitness landscape studies often have to rely heavily on mathematical methods that are not easy to access by biologically trained researchers. Here, we used a method borrowed from engineering - genetic algorithms - to simulate the evolutionary process and study how different components affect the path taken through a phenotypic fitness landscape. In a simple study, we compare five selection models that reflect different degrees of dependency of fitness on trait quality: this includes strengths of selection, trait-quality dependent reproductive hierarchy and the amount of stochasticity in the reproductive process. We include an analysis of other evolutionary variables such as population size and mutation rate. We analyse a game theory problem, as a test landscape, that lends itself to analysis through a deterministic mathematical simulation, which we use for comparison. Our results show that there are differences in the speed with which different models of selection lead to the fitness optimum.

**Author summary:** Evolution and adaptation in biology occurs in *fitness landscapes*, multidimensional spaces representing all possible genotypic or phenotypic combinations, where population adapt by following the cline of the fitness dimension. The study of adaptation on complex fitness landscapes has so far been limited by the need for mathematically heavy methods. Here, we present a simulation modelling framework, genetic algorithms, that can be used for evolutionary simulations of a population on a fitness landscape of chosen features and with custom evolutionary parameters.

## 1. Introduction

### 1.1 Fitness landscapes and evolutionary biology

The evolutionary trajectory of a trait within a population is often represented as a walk in a *fitness landscape* (1) of as many dimensions as the number of (either phenotypic or genetic) components underlying fitness, plus one. The additional dimension is the fitness value that the combination confers to the carrier and is what we use to estimate the evolutionary trajectory of a population. For example, if there are two components then their values can be thought of as orthogonal *x* and *y* coordinates and fitness can be envisaged as the height of a landscape above the *x-y* plan. The shape of the landscape will influence the direction of a notional hill-climber who looks ever to be moving upward. This hill-climbing analogy has been useful in representing and studying evolutionary paths (despite the challenges of visualising a mapping of low and high fitness areas in higher dimensions) and has given rise to several formal models on the correspondence between genotype/phenotype and fitness (2,3). The study of the these models has generated fundamental underpinning to the theory of adaptation, such as the expected distribution of mutation size (4–6). Recent technological developments in genomics and transcriptomics have allowed us to breakdown and reconstruct empirical fitness landscapes (7,8) that are answering our questions regarding their expected characteristics.

So far, no study of evolutionary processes and fitness landscape trajectories has looked at whether varying the relationship between the relative quality of a member of the population and their relative contribution to the next generation affects the evolutionary walk. For example, in a simplified scenario where we measure fitness based on one trait with multiple underlying genotypic or phenotypic components, the same trait variant is likely to achieve different reproductive fitness depending on the reproductive hierarchy within the population. This is likely to influence adaptation in many ecological systems.

The relationship between trait quality and reproductive fitness can be broken down into two key dynamics: 1. the relationship between trait quality and *expected* reproductive fitness (including, specifically in the factors analysed here, the strength of selection acting on the trait and the reproductive hierarchy of the population) and 2. the strength and nature of stochasticity in the reproductive process. Here, we call the interplay of these two factors *selection model* and we study its influence on population-level evolutionary dynamics by means of a simulation method derived from engineering: genetic algorithms.

### 1.2 Genetic algorithms

The exploration *versus* exploitation problem has been explored extensively in a particular subfield of evolutionary computations: genetic algorithms (GAs (9)). GAs were invented in 1960 by John Holland and colleagues as a part of the developing field of evolutionary computations (10). They exploit a natural selection-like process to find optimal solutions to complex problems by evolving a starting population of solutions through selecting, mutating and recombining higher quality variants. This type of heuristic optimisation method is valuable when the solution space analysed is too big to be fully explored and it exploits the correlation in the landscape’s ruggedness. Effectively, it looks for optimal solutions to problems where solutions are given by the non-linear interaction of multiple traits – similarly to a population’s walk in the fitness landscape.

In evolutionary biology, GAs have been applied to problems whose solution by other means might be challenging: searching for efficient parameters for machine learning systems (such as neural networks, *e*.*g*. Montana and Davis, 1989), identifying or evolving solutions to game-theory problems (*e*.*g*. (12,13) and ecological niche models (such as GAs for Rule Set Production, or GARP (14)). They have also been deployed in the simulation of evolutionary or evolution-like dynamics (*e*.*g*. evolution of cognitive processes (15)), although the limitations in understanding and representing analytically what happens during the optimisation process have somewhat limited this latter application.

GAs in evolutionary biology may be usefully applied to investigate how different assumptions about the selection model and, virtually, any other component of the evolutionary process (mutation, sexual or asexual reproduction, structure of the reproductive interaction network within a group, just to name a few) affect the evolutionary trajectory of a trait. Effectively, we can use this tool to extrapolate common rules in the way these components affect the trajectory and to study whether the trajectory holds consistently (or varies consistently) across different fitness landscapes. This knowledge can then inform our prediction of the evolutionary trajectory in natural populations and also improve the theoretical study of evolutionary scenarios. GAs can specifically inform the study of systems where the evolutionary trajectory depends on non-linear (including epistatic) interactions between traits, and thus cannot be captured by deterministic methods (such as adaptive dynamics (16,17).

Here, we start by analysing the effect of the selection model on a small phenotypic fitness landscape that is the product of the interaction of biallelic loci. We assume throughout that reproductive potential across the population is limited to a fixed number of reproductive slots at each generation. We use the five most common models of selection and reproductive slot allocation used in the GA literature as proxies for different biological contexts. These models are: k-tournament, truncation, proportional selection, linear and exponential ranking. We summarise in this introduction how each of these models influences the exploration-exploitation trade-off and we illustrate it with an equivalent biological scenario. We do not claim that our findings obtained by this method can be generalised to all evolutionary adaptive landscapes: as with every simulation modelling method, the identify dynamics that are the result of modelling assumptions – here, the characteristics of the fitness landscape under study. This method allows to the same analysis on different landscapes withing a fixed modelling framework that reduces unwanted sources of variation.

### 1.3 Selection model and the fitness landscape

Recent analyses of empirical landscapes show varying (often considerable) levels of ruggedness, at least for small genomes or gene complexes (18) (with *ruggedness* indicating the amount of epistatic interactions in the genetic architecture (19)). There is, essentially, in biological landscapes a degree of correlation that fits our theoretical understanding of evolutionary landscapes: on average, two neighbouring points in the landscape have similar fitness, while, the more mutations separate them (*i*.*e*. the further away they are in the landscape), the less likely their fitnesses are to be similar, until we reach a distance at which their fitnesses are uncorrelated (19). If the landscape is large enough, multiple fitness peaks might exist far apart. While theory predicts that it is possible (20), if not likely, that a path connecting any two viable genotypes always exist, this path might cross areas of lower fitness, so that a population might find a local peak (a *local optimum*), with higher fitness than the region it started in, but fail to discover the highest peak in the landscape (the *global optimum*). When mutations are not or cannot be large enough to reach distant areas of the landscape, the evolutionary process must rely on strong enough selection to reliably climb fitness peaks, while at the same time allowing exploration of less-fit phenotypes.

Varying selection models might have two distinct consequences on the movement of a population in the fitness landscape and we illustrate them here. Let us take a classic case study evolutionary scenario used in the evolutionary literature: a discrete fitness landscape with neighbouring points assumed to be at one-mutation distance from each other and with mutations assumed to be rare enough to arise one at a time. If the evolutionary process relies on a small group of high-quality individuals to reproduce, then the population will follow a highest-fitness neighbour trajectory and is likely to land on the closest fitness peak in the landscape. If this point is a local optimum, that optimum is where the population will stay until the next change in selective pressure that modifies the landscape. Maintaining sufficient variation in the population, by allowing lower quality individuals to reproduce, on the contrary facilitates the emergence of new variants away from the current optimum. A second consequence of the relationship between trait quality and reproductive fitness is the change in the speed with which the population moves along the trajectory: the higher the proportion of reproductive fitness assigned to the highest-quality variants, the quicker the movement. The higher the proportion that lower-quality variants benefit from, the more time the population needs to move to a point with higher average quality.

### 1.4 Explorative and exploitative selection models

We define *exploitative* systems as those biological scenarios where the highest-quality individual (or a small élite of high quality individuals) obtains a share of reproductive fitness that is disproportionately large compared to what would be expected should reproduction be directly proportional to the relative quality of the individual with respect to the average. An exploitative system is, for example, a non-eusocial group with a single reproductive male or female which suppresses subordinate reproduction and sires all new offspring. Conversely, an *explorative* system corresponds to a scenario where most individuals have at least as much reproductive fitness as what would be expected from their relative quality in the population. In the absence of sexual selection (or in the presence of moderate sexual selection), biological scenarios in which individuals compete for each reproductive opportunity (for example, species in which reproductive pairs form and dissolve yearly) can be considered explorative and the amount of exploration is increased by stochastic events interfering with the fitness-proportional distribution of reproductive chances (*e*.*g*. in a species with a 20 year reproductive span, the individual with highest fitness expectations might stochastically die after only 1 year, but the individual with the lowest expectations could, stochastically, reproduce for 11 years).

In the study described below, we investigate how different systems affect the evolutionary walk of a population in a phenotypic landscape from game theory, the SPS game.

### 1.5 Landscape of the Sir Philip Sidney game

The fitness landscape we study here, the Sir Philip Sydney (SPS) game theory problem (21), has been chosen for ease of access and understanding at the time this study began and it cannot be described purely in terms of well-established formal models of fitness landscapes. Nevertheless, those shortcomings do not prevent it from being a useful representation of a biological landscape. It is a phenotypic gambit scenario of six independent traits contributing to the total reproductive fitness of the individual. Mutations are additive at the phenotypic level, but non-additive at the fitness level, meaning that the traits interact non-linearly to determine fitness. Our landscape differs from the assumptions commonly made in the literature in two ways. Firstly, it is discrete. Mathematically, this is the equivalent of any continuous landscape whose optima and minima are all represented on our discrete landscape. Secondly, it is a multi-phenotype frequency-dependent landscape. Although this is not a commonly studied landscape scenario, it is found in several real-world evolutionary case studies, such as apostatic selection or microbial competition experiments where multiple strains grow on a medium with multiple carbon sources.

### 1.6 Details of analysed selection models

We call selection model the set of assumptions in an evolutionary simulation that define the relationship between trait quality and reproductive success of the individual, as a function of the two dynamics listed in paragraph 1.1 (relationship to expected reproductive fitness and amount of stochasticity in the system). Different biological scenarios often imply different selection models. Below, we provide a brief summary of the models analysed here.

#### k-tournament

This selection model assumes that competition for each reproductive slot occurs in small, randomly formed groups, where the individual with the highest-quality trait wins with a specified probability. The stochasticity introduced by randomly selecting competitors and the uncertainty in the probability that the individual with the highest-quality phenotype wins means that *k*-tournament allows for some stochasticity in the selection of the traits transmitted to the next generation. This method is equivalent, in a biological scenario, to a group of individuals competing for a mate after being drawn to the same location by a mating signal: competitors are brought together by contingency and the strongest does not necessarily, albeit is more likely to, prevail.

#### Truncation

Truncation sets a fixed threshold to the proportion of the population that can reproduce and makes this threshold dependent on trait quality. For example, only the top 10% of individuals in terms of trait quality may have the opportunity to reproduce. Among this reproductive élite, chances of reproducing are then uniform (22). Making a parallel with a biological scenario, we can compare this method of selection to dynamics in an animal group where a reproductive hierarchy is established, by which only top-ranking individuals mate and reproduce.

#### Proportional selection

Proportional selection allocates reproductive slots in a manner that is statistically proportional to trait quality, but it introduces some stochasticity in the allocation. In a biological equivalent, we can think of this method of selecting reproductives as a system where individual reproductive success is strongly dependent on the absolute quality of one trait – but failure of a high-quality individual to reproduce is still possible (simply through bad luck).

#### Ranking

We can visualise the ranking selection model as a population organised as a list of individuals ranked from highest to lowest trait quality. Individuals are then assigned reproductive slots according to the place held in the ranking (plus some additional stochasticity), rather than to the absolute trait quality value. The advantage of this method is that the within-population variance in the expected number of reproductive slots remains fixed, evening out reproductive chances within each generation and effectively enhancing exploration. In fact, because the range of the number of allocated slots is independent of population trait-quality mean and variance at that time, the highest quality phenotype in the population will receive the same expected reproductive proportion regardless of how far away it stands from average population trait quality. Conversely, lower quality individuals will enjoy reproductive opportunities as long as their ranking position is not too far below the rest of the population. A biological equivalent of this method is a scenario with a hierarchically organised population where reproductive opportunities are proportional to the rank and where the number of reproductive positions available remains relatively fixed across generations.

We also use this study to test the effect of other evolutionary parameters on the evolutionary trajectory. The complete list of evolutionary parameters tested (including selection method parameters) is given in **Table 1**. We explain here the use of one particular method, derived from GA literature, and its meaning as a biological parameter: *replacement*.

**Table 1.**
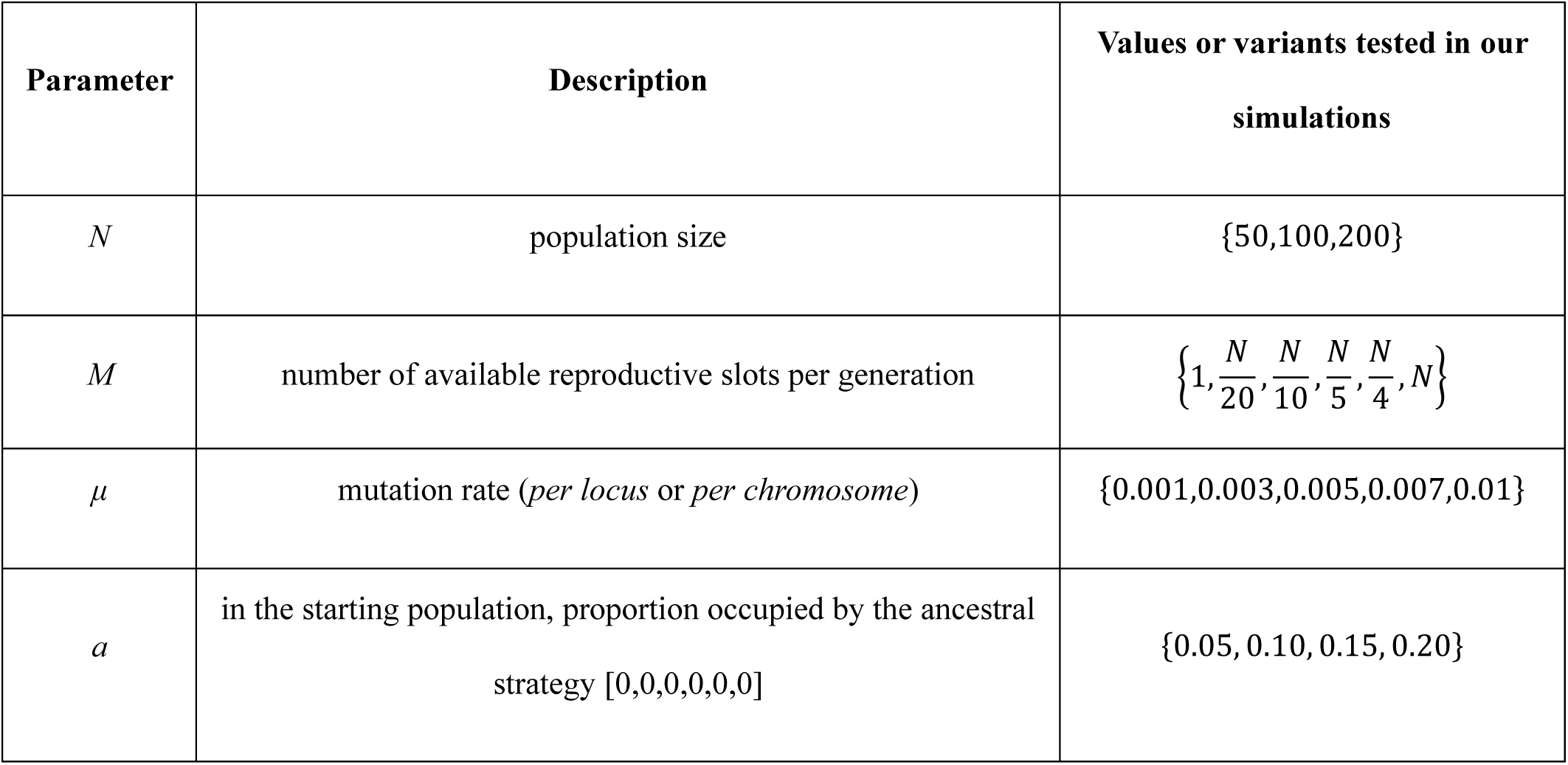
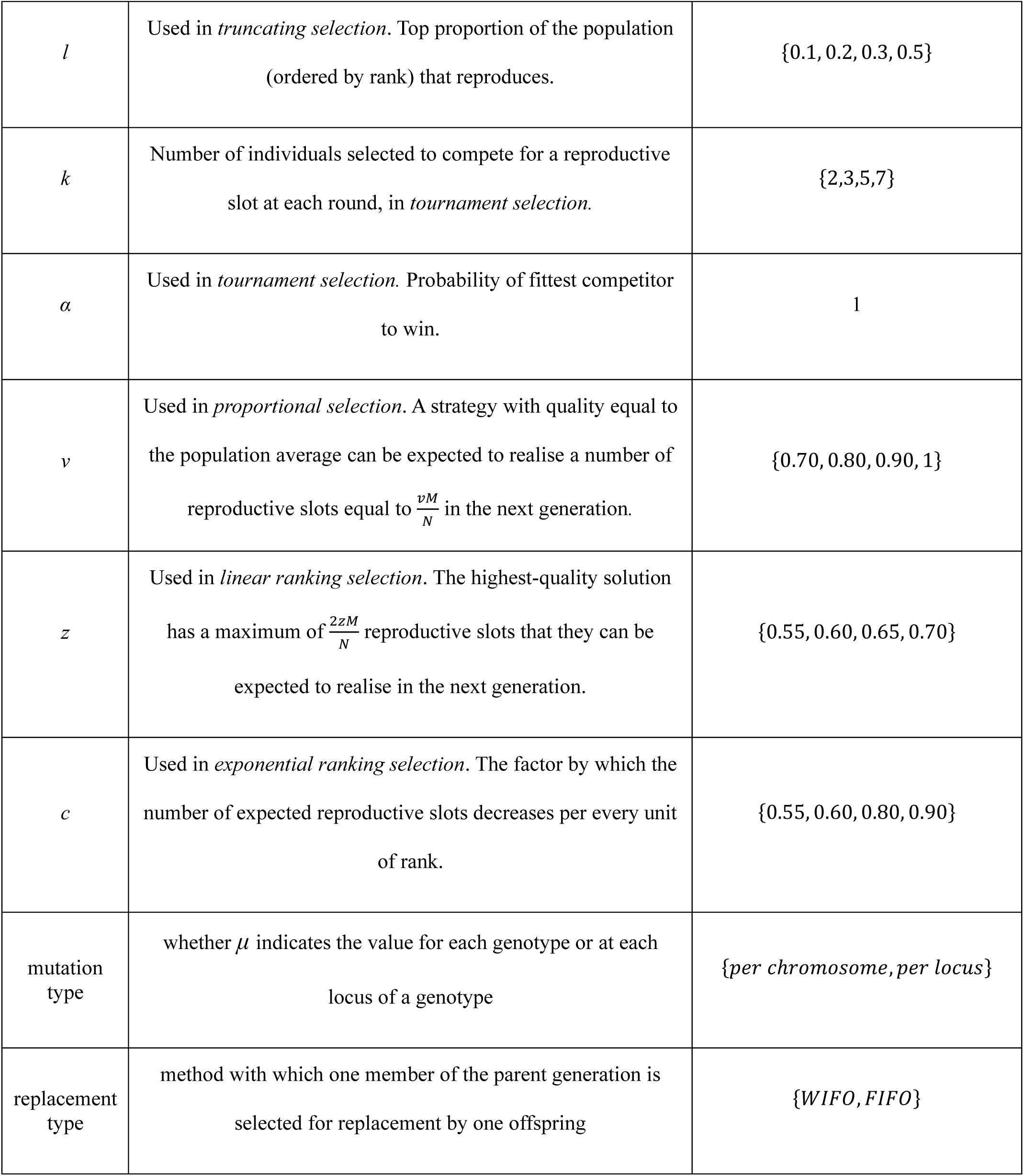
Description of parameter tested with listed values.

In a GA, replacement is the process by which new individuals replace members of the parent generation in a fixed-size population. It relies on two features with important evolutionary implications in affecting the exploration-exploitation balance: replacement size (or number of reproductive slots *M* relative to population size, *M*:*N*) and replacement criterion.

It is either generational, where the offspring completely replaces parents in the new generation (*M* = *N*), or overlapping, where only a fixed proportion of the population is replaced (*M*<*N*). In biological terms, this is the equivalent of studying the effect of the reproductive overlap between generations. Overlapping replacement allows trait variants already present in the previous population to compete with newly-generated variants. Increasing *M* considerably reduces variance in the growth curve of the best-variant proportion of the population, with generational replacement having the least variance (23). It thus reliably produces quality-proportional strategy fitness, but relative to the quality of already-existing strategies – the local fitness cline is exploited. The second critical feature is how strategies are selected for replacement. In the First-In-First-Out (FIFO) GA method, individuals are replaced in chronological “birth” order, while in Worst-In-First-Out (WIFO) replacement occurs either by quality-dependent proportional selection or by deterministic death of the worst individual. This is the equivalent of studying the effect of the relationship (or absence thereof) between trait quality and probability of death on the evolutionary trajectory. FIFO favours drift and thus exploration, granting each strategy the same reproductive time frame, while WIFO is exploitation-based.

In this paper, we describe the results obtained from evolving a population towards a solution to the SPS game, under each selection model and across different parameter values for selection, mutation, population size and replacement. Multiple solutions to the SPS game have been found by different methods (see **Box 2**) and are considered the expected endpoints of an evolving population. We compare the reliability with which each model reaches the evolutionary stable solutions previously identified for this game: the evolutionary stable strategy (ESS) analytically found by Johnstone and Grafen (24) and the evolutionary stable set (ES) obtained by Hamblin and Hurd (25).

### Box 1: Individual, genotype and phenotype in a GA and in biology

In a GA, populations consist of multiple possible solutions to the problem under study that are undergoing a selection process for a high-quality solution. This population is the equivalent of a biological population of individuals displaying variation in the trait (solution) under selection. Each solution is encoded by a genotype, consists of a string of values, one (*locus*) for each of the parameters or features that contribute to a solution. For example, in a GA genotype modelling leopard predatory effectiveness as a physiological trait, the genotype might consist in a series of loci encoding the parameter values for: muscular mass, fibre elasticity and the parameter defining the developmental and physiological trade-off between the two. The solution built under this genotype is a combination of muscular power and elasticity used to respond to predatory encounters, in which prey size and speed are drawn from fixed probability distributions. While phenotype and genotype overlap in one-locus traits, the difference becomes apparent wherever we introduce an interaction between loci. In our predatory effectiveness trait, it is the trade-off parameter that causes a phenotype to be different than the effect of the power and elasticity loci taken independently. This resembles the complexity observed in real biological traits. Similarly to biology, moreover, an additional layer of complexity can be introduced at the strategy-generation step if noise or plasticity (learning) mechanisms influence the trait. Multiple individuals in the same population might carry the same genotype and, indeed, the change in genotype frequency in a population over generations is one currency for measuring its success.

In the game theory problem modelled here, we are looking at the evolution of a behavioural strategy, in a signalling-for-resources scenario. Each strategy is a combination of behavioural responses, each encoded at a separate locus. Here, we call the behavioural strategy a *strategy* (*solution*) and we follow GA terminology in using *genotype* to indicate the combination of behavioural responses that make up a strategy, despite them being the phenotypes in the phenotypic landscape.

## 2. Methods

To test the effect of selection models over evolutionary trajectories, we evolved solutions to the Sir Philip Sidney game under different models, using a genetic algorithm. We applied five different selection models, each evaluated across a range of parameter values. **Table 1** summarises the parameters analysed and lists the values tested for each.

### 2.1. Genetic Algorithm

The workflow of a GA is depicted in **Figure 1**.

**Fig 1.**
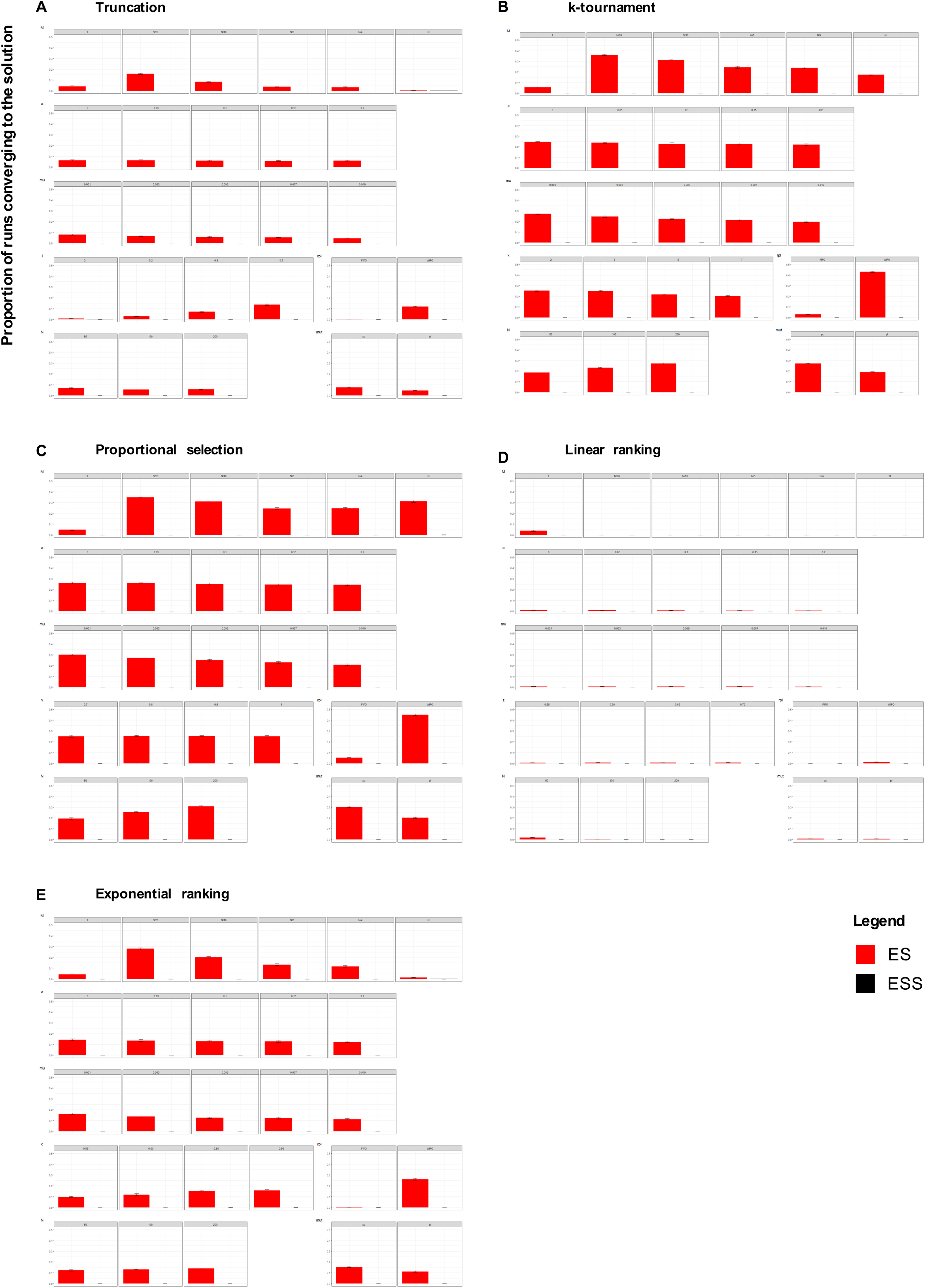
Workflow of a GA. Simplified representation of the processes in a GA (in white), that pool, select and modify the initial *population* (in red) to generate a new population. The *parental pool* are the individuals in the current population that meet the trait-quality requirements to reproduce. They produce copies of themselves (*parental copies*) that undergo mutation and crossover to generate the final *offspring* individual, as in asexual selection. Offspring take the place of individuals of their parent’s generation in the population. Each cycle through this sequence of steps constitutes one generation.

#### 2.1.1. Chromosomes

Strategies are encoded by 6-locus strings, or *chromosomes*, with each locus encoding either a 0 or a 1.

#### 2.1.2. Population and replacement size

The population consists of *N* individuals, of which *M* are replaced by offspring strategies at each generation.

#### 2.1.3. Trait quality evaluation

This is where the quality of the strategy generated by each individual *n* with genotype g_*i*_ in the population is tested. We define trait quality as the average outcome of the test; this value is converted into fitness at later stages. In this simulation, quality is evaluated through competition against randomly selected strategies from the same population over five rounds, as in (25). At each round, an “opponent” strategy is chosen among the *N* – 1 remaining and takes the complimentary role (*i*.*e*. donor or signaller) to the strategy evaluated. Role, health state and degree of relatedness are assigned with probability equal to population frequencies in the equilibrium parameter range of (24). Performance is calculated from the interaction of the two strategies in each round and trait quality as the average of all rounds.

#### 2.1.4. Selection

We assume throughout that reproduction is asexual, requiring only a single parent. Selection defines which subset of the parental population will be passed on to the next generation and how many breeding slots (proportion of the offspring generation) each parent solution receives.

##### k-tournament

*k* randomly selected individuals are compared in each of *N* rounds. In each round, the individual with highest quality is selected with probability *α*. If the highest-quality individual is rejected, then the second fittest is chosen with probability *α, etc*. Selected individuals re-enter the population pool after reproduction and can thus be selected multiple times.

##### Truncation

Individuals are sorted by decreasing quality and parents selected from the *l* topmost proportion of the list are selected for reproduction. Note that individuals with equal trait quality might be separated by the threshold and some not enter the selection pool. Parent individuals are selected with uniform probability from the selection pool to enter the parent pool.

##### Proportional selection

Individuals are assigned an expected value of reproductive slots, which is, the statistical mean of the number they should receive in an infinite number of trials. The assignment function used to derive expected values is

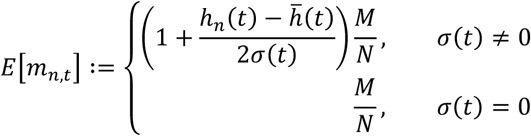

from (26). The equation represents the expected number of reproductive slots *m*_*n,t*_ that individual *n* will be allocated at time *t*, given its estimated quality at that time step, *h*_*n,t*_, and the average quality in the population at that time step, 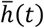, and adjusted according to the standard deviation of quality within the population at that time step, *σ(t)*. Variance-scaling is applied to limit the effect of drift from the highest quality value, making reproductive chances more widely distributed when variance is high and less when it is low. We modify (26) to control how reproductive slot assignment depends on the quality value. Via proportional selection, we can set the expected value for an individual with average quality through a parameter *v*. This is done by multiplying:

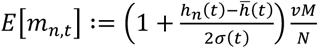

When *v* =*1*, an average quality individual receives as many reproductive slots as if assigned by a uniform quality-independent distribution. If we set *v<1*, only higher-than-average individuals will expect at least one reproductive slot. The lower *v*, the greater the reproductive advantage of high-quality individuals over those of average quality. Thus, the lower *v*, the higher the exploitation.

The effective number of slots is assigned using a sampling algorithm: a random integer *r* between 0 and the sum of expected values is drawn for a number of rounds *M* equal to the number of reproductive slots and the expected value of individuals summed as the population is looped through in a fixed order, until the expected value of an individual makes the total equal to or higher than *r*and that individual wins one reproductive slot. Slot allocation following this system is thus statistically proportional to assigned expected values, with drift. *Linear ranking*: Individuals are ranked in increasing order of quality and the rank used to assign the expected value through an assignment function. The linear implementation of the assignment function (27) is:

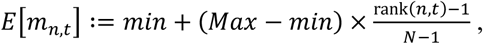

where *min* is the expected value allocated to the worst individual in the population at time *t* and *Max* that allocated to the best individual. In this type of implementation, it is possible to manipulate the expected fitness of the best individual to adjust the exploitation-exploration balance, in the same way as we changed the expected fitness of the average individual in proportional selection. *Max* can be set to a different value from its maximum, *2M/N*, through a parameter *z* similar to *v* in proportional selection, so that

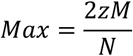

(see the derivation in SI.1.).

The effect of *z* is analogous to that of *v* in proportional selection, but with opposite effects as it acts on the highest-quality individual: the higher *z*, the higher the exploitation and the lower *z*, the higher the expected fitness of lower quality individuals.

Reproductive slots are then assigned with a sampling algorithm identical to that in proportional selection.

##### Exponential ranking

The exponential implementation of the assignment function takes the form

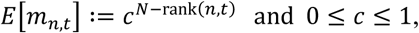

where *c* determines the steepness of the exponential increase in expected values with rank. By changing *c*, we can control the exploration-exploitation balance: a high *c* assigns exponentially higher advantage to the highest-quality individual, thus creating high exploitation.

The effect of *z, v* or *c* is that of fine-tuning the proportion of individuals effectively reproducing (what Baker (27) calls “percentage involvement”).

#### 2.1.5. Mutation

Parents that are assigned breeding slots through selection generate copies of themselves. Such copies undergo mutation to generate new variants and explore the solution space around parent solutions. We do not explicitly assume a “strong selection, weak mutation” (SSWM) regime: we instead model mutation through a mutation rate parameter and a mutation method parameter that span through a range of mutation regimes. In *per chromosome* mutation, one locus on the chromosome is selected with uniform probability and with probability *μ*_*c*_its value switches to the opposite binary value. In *per locus* mutation, the chromosome is scanned and every locus switches with probability *μ*_*l*_. As a result, our study investigates SSWM conditions, under low mutation rate, per chromosome method and small reproductive slot size, as well as biologically unlikely high-mutation scenarios, when reproductive slot size is very large or under per locus mutation. We chose not to include crossover in the analysis, because, in small fitness landscapes, mutation should be sufficient to explore the full space. In accordance with this expectation, (25) found that crossover does not affect the endpoint reached. After mutation (and crossover), the offspring set has been generated and is input into the population, entering the next generation.

#### 2.1.6 Replacement

At the end of each cycle, *M* individuals among the population’s *N* are replaced by the offspring. Individuals in the parent generation are chosen for replacement either on the basis of quality (Worst-In-First-Out or WIFO), where the *M* lowest quality individuals are replaced, or in a time-sequence manner, with oldest individuals being replaced (First-In-First-Out or FIFO). We investigate both WIFO and FIFO.

#### 2.1.7. Running time and data collection

The algorithm is run for 500 generations, as in (25). Frequency of each possible strategy in the population and population fitness values are recorded every 50 generations.

#### 2.1.8. Implementation

Each combination of selection model and parameter values was run in 10 replicates, each with random number generators seeded with a different integer value between 0-9. The simulations were implemented in Python version 2.7.12 using Cython language version 0.27.3.

### Box 2: Sir Philip Sidney game

The SPS game is a game theory problem in evolutionary biology used (first by (21)) to study the evolution of costly and honest signalling within an inclusive fitness framework. The problem of signalling for help when health is poor is evolutionarily interesting because, once honest signalling has evolved, individuals who are in good health benefit by signalling as if they were poorly and receiving the resource at the expense of the donor. This means that honest signalling is evolutionarily unstable. However, honest signalling might be maintained through a combination of inclusive fitness - conferring an indirect fitness benefit to the helping donor and to the honest signaller - and of signals that are costly to produce - enforcing lower dishonesty. The SPS game recreates a simple case study scenario with a population of individuals of relatedness *r* who can transfer a fitness-increasing resource to each other. Individuals can choose whether to ask for the resource, paying a cost *c* to signal for help, and whether to donate it, lowering their fitness but increasing that of the receiver. Individuals should take into account their own level of fitness (usually called “thirst”, for fidelity to the Sir Philip anecdote), as well as the potential donor’s, to decide whether or not to signal. The game asks which conditions of relatedness and cost allow the evolution of completely or partially honest signalling.

The mathematical models following Maynard Smith’s (*e*.*g*. (24,28–31)) have tested the problem under different scenarios by varying underlying assumptions; however, evolutionary simulations have challenged the notion that ESSs, obtained through either mathematical analysis or deterministic evolutionary simulations (*e*.*g*. (32)) and albeit possible under specific conditions, should be the expected evolutionary outcome.

### 2.2. SPS game

We use the SPS game (see **Box 2**) as a testbed problem. We chose to reproduce the study by Hamblin and Hurd (25), who evolved solutions to the SPS scenario investigated by Johnstone and Grafen (24) using a GA. The signalling scenario studied by (24) consists of a population of donors and of two types of signallers, a close relative with relatedness *r*_*1*_ to the donor (type I) and a distant relative (type II) with relatedness *r*_*2*_. Signallers can be in either of two states: thirsty and dying in the absence of the resource (fitness = 0), and healthy, with fitness *F*_*b*_ in the absence of the resource. Resource transfer re-establishes full fitness. Signallers are healthy with probability *o* and needy with probability (1 - *o*) and closely related to the potential donor with probability *q* and distantly related with probability (1 – *q*). Similarly, donors have fitness of 1 with the resource and of *F*_*d*_ without. The donor can decide whether or not to transfer the resource, based on whether a signal received, and signallers can decide whether to signal or not. Signalling has a cost *d* = *d*_1_ ⨯ F for close relatives and of *d* = *d*_2_⨯ F for distant relatives, where *F* is the signaller’s fitness after the donor has made its decision. The optimal strategy for the donor will depend on the degree of relatedness to the signaller and on the proportions of relatives of each degree in the population, while the optimal strategy for signallers will depend on relatedness and the cost of signalling.

We searched for solutions for the same SPS game parameter set used by (25) and based on (24) (also accepting (25)’s correction for *d*_*2* =_ 0.1 for the semi-separating equilibrium to be an analytical ESS solution to the game). This parameter set is a point in parameter space at which the analytical ESS is expected. This ESS is a biologically interesting case of partially-honest signalling and is the highest-payoff strategy when only one strategy is present in the population (at this point in parameter space). However, most commonly the population seems to reach a different convergence point, when an evolutionary simulation is used (25): the ES, a two-strategy endpoint that has the highest average payoff in the population when every strategy is equally represented at time zero (see **Supporting File 1**).

#### 2.2.1. Chromosome representation of a strategy

The six loci represent, in order: donor strategy on signal received, donor strategy on no signal (1 = transfer resource, 0 = no transfer), closely-related signaller strategy when thirsty, closely-related healthy signaller strategy when healthy, loosely-related signaller strategy when thirsty, loosely-related healthy signaller strategy when healthy (1 = signal, 0 = no signal) (**Figure 2a**).

**Fig 2.**
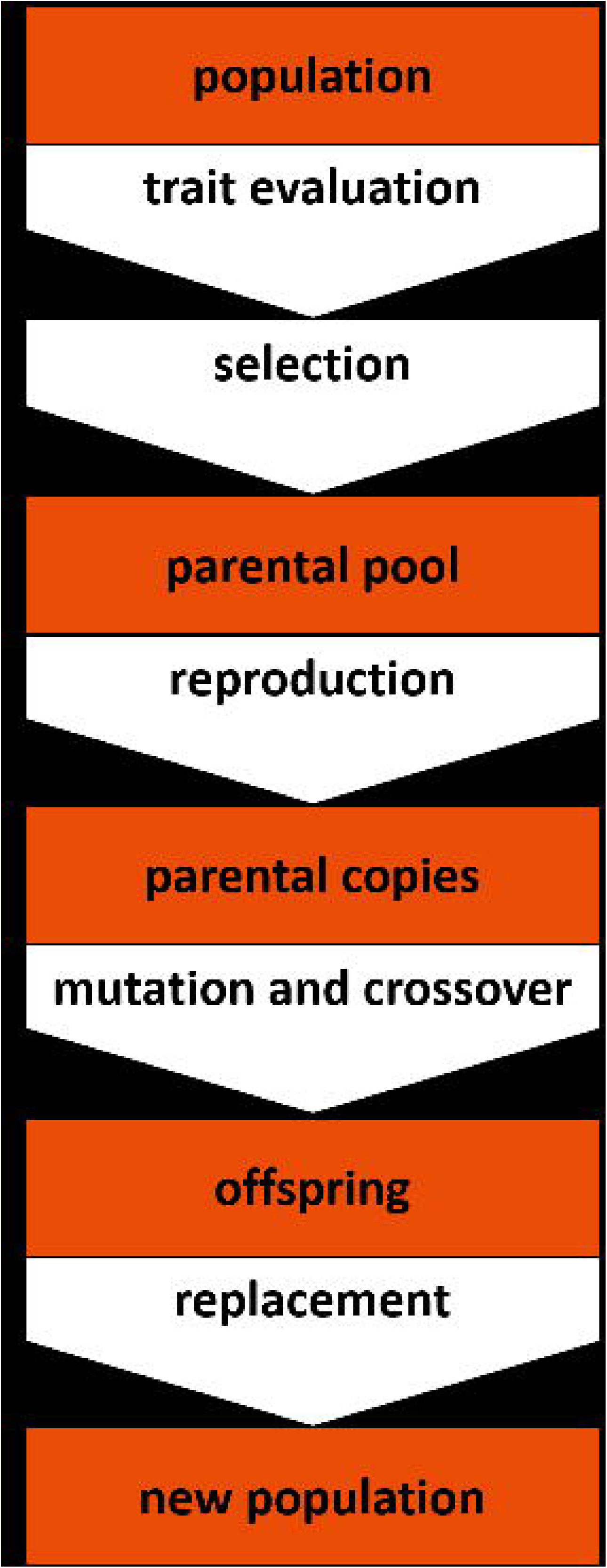
How SPS strategies are encoded in the GA. **A**. Structure of a GA chromosome encoding a strategy used in the SPS game. Each locus (A-F) contains a binary value indicating whether that behavioural response is used by the carrier (1 = yes, 0 = no). The first two loci are behavioural responses from the donor of the resource: transfer the resource when hearing the signal (A) and transfer the resource when *not* hearing the signal (B). The other four loci encode the responses of the signallers to their own health status and to the degree of relatedness to the donor. Loci C-D encode the responses towards closely related donors: emit a signal if thirsty (C) and emit a signal if healthy (D). Loci E-F encode the same responses towards distantly related donors. **B**. The table shows how the ESS and the two strategies within the ES are encoded according to the scheme above.

### 2.2.2. Initialisation

At the start of each simulation, a proportion *a* of the population consists of the putative ancestral strategy [*no-transfer, no-signal*]. The remaining strategies are randomly drawn among all possible SPS game solutions.

#### 2.2.3. SPS game parameter values

Parameter values used: *o* = 0.6, *q* = 0.9, *r*_*1*_ = 0.5, *r*_*2*_ = 0.2, *F*_*b*_ = *F*_*d*_ = 0.8, *d*_*1*_ = 0.4, *d*_*2* =_ 0.1.

At this point in parameter space, we find the ESS and the ES detailed in **Figure 2b**.

### 2.3. Deterministic model

To have a benchmark for estimating the amount of exploration generated by each selection model, we produced a deterministic model and compared its results with those obtained by the GA. The model assumes a population of fixed size equal to the total possible strategies in the SPS game (*N* = 64). At generation 0, each strategy is equally represented in the population (*i*.*e*. there is exactly one copy of each strategy). Fitness in the model is given exclusively by the relative quality of each strategy. We define the fitness of strategy *i* at time *t* as

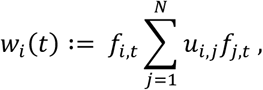

where *f*_*i,t*_ is the frequency of strategy *i* in the population at time *t* and *u*_*ij*_is given by the average value obtained by *i* when competing against strategy *j* in the SPS game, defined as the average between playing as the donor and playing as the beneficiary. These two components of *h*_*ij*_were calculated as in (24), for the same values of SPS game parameters used in the GA (*i*.*e*. for the point in the SPS parameter space where the ESS is the strategy detailed in **Figure 2**). The average fitness obtained by all strategies at time *t* is

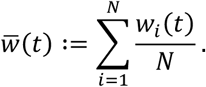

The relative fitness of strategy *i* is thus

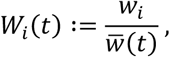

which becomes the frequency of strategy *i* at time *t+*1.

The model has no mutation nor recombination. We calculate population frequencies for 500 generations, which is the same number of generations used in the GA.

The deterministic model was built and run in MATLAB R2020a. The matrix of values resulting from the interaction of all pairs of strategies was built in Python version 2.7.16.

### 2.4. Analysis

#### 2.4.1. Fitness landscape exploration

To quantify the effect of parameters and methods on landscape exploration, we only considered those simulations in which the ESS or ES reached and maintained a population-level frequency higher than 80% for the last 50 generations of the simulation. We consider those runs as having converged to that solution (as in (25)). For each selection model, we calculated the proportion of runs that converged to each solution type. Within each selection model, we then compared the effect of each parameter value of each evolutionary parameter by looking at the proportion of simulations with that value that had reached each solution type, across all values of other parameters. Mean and standard deviation were calculated as the mean and standard deviation of replicates with the same seed. In total, 72,000 simulations were run for each selection model and 12,000-36,000 simulations were run for each parameter value within selection model (that is, across all values of other parameters). This difference in the number of simulations for each parameter value is an effect of grouping runs along single parameter dimensions: for example, if we calculate statistics for each value of *M* within one selection model, we analyse 72,000/6 = 12,000 simulations (where 6 is the number of values of *M* we tested in total; see **Table 1**), while, if we do the same calculations for *N*, we look at 72,000/3 = 24,000 simulations.

#### 2.4.2. Statistical analysis

We tested for a significant difference in the distribution of number of seeds converged, across selection methods, to three solution types: ESS, ES and all other solutions. We used a Fisher’s Exact test with simulated p-values by Monte Carlo simulations, 10,000 iterations. Statistics were implemented in R, version 3.5.1 (33).

## 3. Results

### 3.1. Different selection models lead to different outcomes

We reproduced the analysis run by Hamblin and Hurd (25), who used a GA to explore a point in the parameter space of the SPS scenario studied by Johnstone and Grafen (24). We used this case study to analyse the effect that different assumptions made on the evolutionary process have on the trajectory of a simulated evolving population. Specifically, we were interested in the evolutionary outcome under multiple selection models. We also looked at the effect of the evolutionary parameter values listed in **Table 1**. Hamblin and Hurd’s algorithm reaches quite different evolutionary endpoints from the original 1993 study of this SPS game variant, with the ESS identified analytically at this point in parameter space ([*give-on-signal, signal-when-needy, always-signal*] for [donors, closely-related signallers, loosely-related signallers]) effectively emerging with very low frequency. More frequent as a solution is an ES of strategies, [*give-when-no-signal, never-signal, never-signal*] and [*always-give, never-signal, never-signal*].

We analyse the effect of each selection model on the point in the phenotypic landscape which a population reaches within 500 generations. We compare these results to the convergence point of a fully deterministic (*i*.*e*. perfectly fitness-proportional) model run for the same number of generations, to better understand the amount and effect of exploration in each of our partially stochastic selection models. For convenience, we define three types of solutions to the SPS game: the Evolutionary Stable Strategy (ESS), the Evolutionary stable Set (ES) and every other solution. We can envisage that exploitation of the best individual will lead to the ES, more commonly than to the ESS, due to the former being the highest *average* fitness strategy, while the ESS needs high frequency in the population *before* reaching the highest payoff. In accord with this latter statement, we can see in (25) that ESS convergence increases with the frequency at which it is seeded in the initial population.

If we look at the proportion of simulations reaching the two evolutionary stable solutions (ESS and ES) in our study (**Figure 3**), we see that no model reliably reaches the ESS and that there is no difference among the models in the frequency with which they land on this solution (**Figure 3a, bottom panel**). The ESS is thus not an evolutionary attractor. When we shift our attention to the ES, we realize that different selection models reach the highest fitness solution with very different frequencies (**Figure 3a, top panel; note the difference in the y-axis scale with the bottom panel**; Fisher’s Exact test, p-value<0.001).

**Fig 3.**
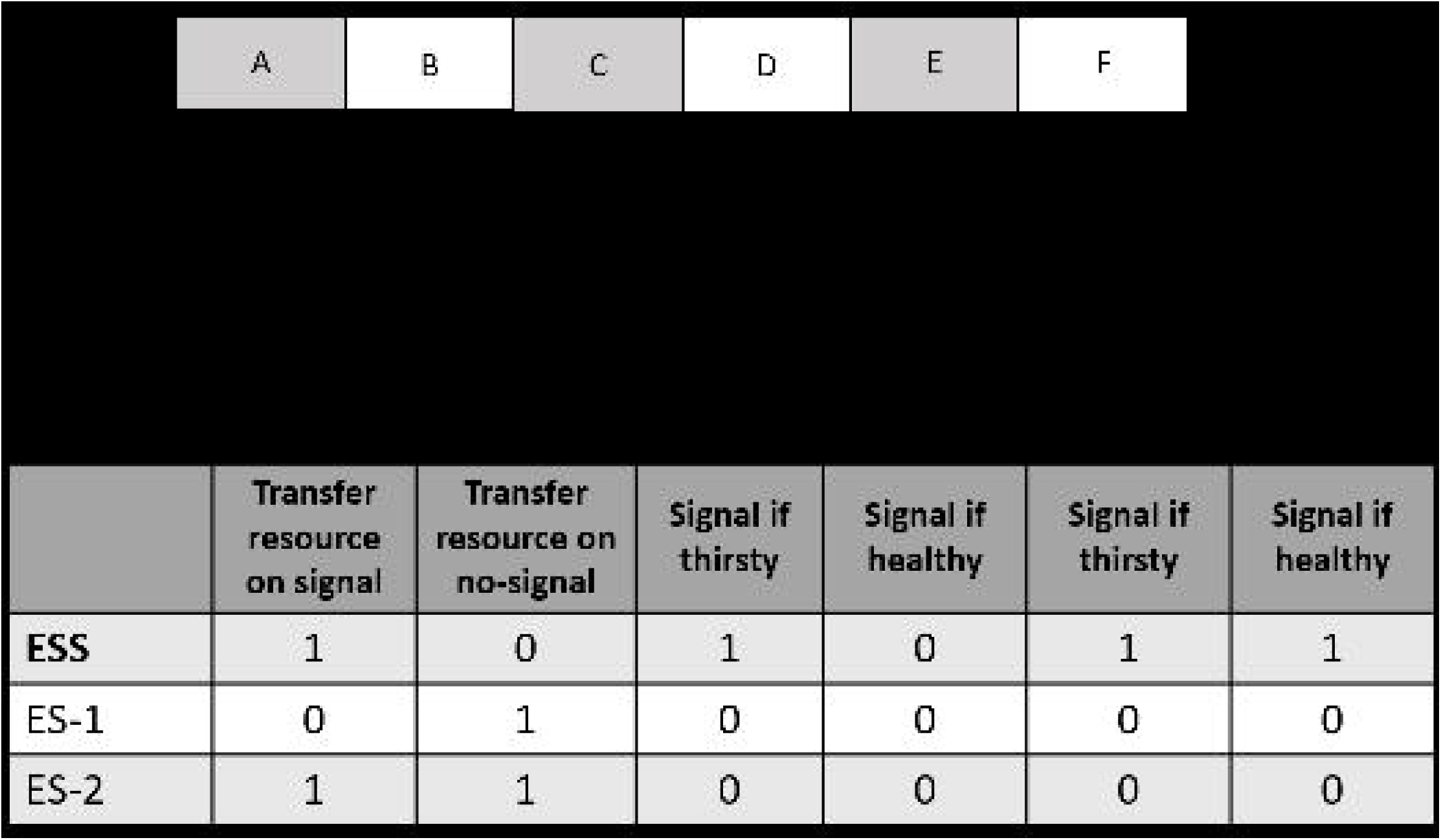
Proportion of simulation converging to ESS or ES by selection method and deterministic fitness of ESS and ES. **A**. Proportion of simulations that converge to each solution type, of those run with the specified selection method (across all other parameters and parameter values). Note that the top and bottom panel are plotted across different y-axis values. The standard deviation bars reflect the variation among runs with different random seeds, collapsed across all parameter values. **B**. In the deterministic model, the ES and the ESS reach these frequencies at generation 500. The frequency of the ESS is 3.8e^-47^. These frequencies are the outcome of a selection process where fitness exactly corresponds to relative trait quality within the population, with each strategy being present once in the population at time zero and in the absence of mutation.

Most of our selection models reach the ES with very high frequency. If we consider that, in a selection model where fitness equals relative trait quality such as the deterministic model, the ES only reaches a frequency of 0.5 in the population in 500 generations (**Figure 3b**) and that we have set the threshold frequency to consider a strategy as the evolutionary endpoint to 0.8 (maintained over the last 50 generations), these frequencies are considerably higher than expected. What is happening here? Reproductive fitness in our model is dependent, as in most biological scenarios, on an estimate of trait quality. This estimate is obtained from five “events”, each under randomly assigned “conditions” (relatedness, health state, identity of the other strategy and the role played, in this SPS game) – it is a combination of stochasticity and of *median* quality, rather than average, that creates the estimate. Our ES has high estimated quality under most condition (**Supplementary file 1**: strategy 1 in the ES has the highest payoff in almost half of the pairwise competitions with other strategies: 30/64, including itself; strategy 2 has the highest payoff only in 3/64 pairwise competitions, including against strategy 1 and itself – it is an equivalent phenotype to strategy 1 at equilibrium). However, many other strategies have high quality in more than one condition. In many conditions where the ES has maximum quality, one or more other strategies share this property. Essentially, the average landscape is relatively flat, which means that there is a good chance of error when estimating overall trait quality from only a small number of samples.

Competition within small groups (k-tournament) favours strategies with high mean estimated fitness: an individual carrying that strategy has low chances of competing against an individual with higher estimated fitness and, if they win, they automatically reproduce. Larger sizes of competition groups slow down the diffusion of the ES (**Figure 4b**). Even in a relatively flat landscape, assigning fitness in a manner directly proportional to estimated trait quality is enough to reach quick and reliable convergence to the ES (proportional selection). This is independent of the estimated number of reproductive slots assigned to the average individual, *v*, (**Figure 4c**) and we believe this is because drift occurs only towards high quality, not average, solutions (the average mean payoff from pairwise interactions among all possible strategies is 1.056; the average mean payoff of strategies that obtain the highest payoff against at least one other strategy is 1.095). Ensuring that everyone in the population reproduces (ranking selection) makes convergence slower. Even a reproductive hierarchy that assigns a lot of advantage with every step up in the rank (exponential ranking) reaches the ES in 500 generations with much lower frequency than with competition within groups (k-tournament) or trait-quality proportional selection. Increasing the advantage for each step in the rank (*c*; **Figure 4e**) increases the speed of convergence. Linear ranking is the slowest method and seems to work at its best in small populations (*N*; **Figure 4d**). However, limiting reproduction to the individuals with the highest estimated quality (truncation) also slows down convergence to the ES. The larger the share of the population that is allowed to reproduce, the faster the convergence (*l*; **Figure 4a**). Based on these observations, we think that a strict regime of selective reproductive allocation accentuates the effect of the error in the quality estimate.

**Fig 4.**
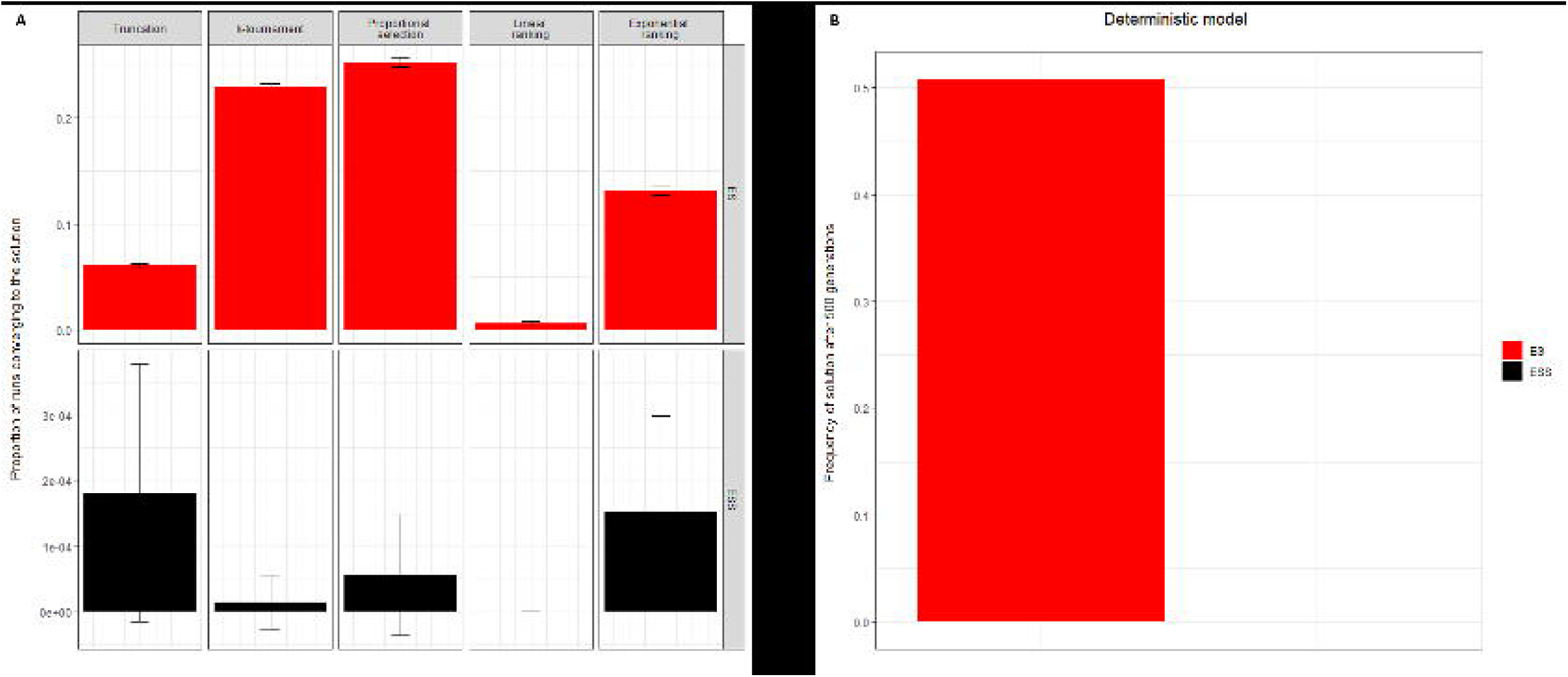
Proportion of simulations converging to ESS or ES by parameter value, within each selection method. Each group of plots shows the proportion of simulations that converge to each solution type by parameter type and value (*i*.*e*. across all other values and all other parameters), within a selection method. The standard deviation bars indicate the variation found among runs with different random seeds, collapsed across all parameter values. mut = mutation, pc = per chromosome, pl = per locus, rpl = replacement.

### 3.2. Effect of other parameters and methods

We can identify some general effects of other evolutionary parameters and of the mutation and replacement processes by looking at the general performance across selection models (**Figure 4**).

- In this small fitness landscape, a high mutation regime decreases the speed and reliability of convergence. However, even under very high mutation rate (*µ*) values, most methods reliably reach the ES in 500 generations.
- A trait-quality dependent death rate (WIFO) increases the speed of convergence across all selection models, compared to an age-dependent death rate (FIFO) (this is in agreement with results found in GA literature (34)).
- A larger population size increases convergence. The effect seems to level off once population size becomes sufficiently larger than the solution space (>64), although we would need to run simulations with intermediate population sizes (*e*.*g*. 30<*N*<100) to confirm this effect. The effect of large *N* on convergence is true under all selection models, with two exceptions. Under linear ranking, a small population size increases the frequency of convergence; under truncation, population size seems irrelevant – probably because the key parameter here is the proportion of the population that is reproductively active, *l*.
- Having an intermediate number of reproductive slots available at each generation accelerates convergence, probably by counterbalancing the high mutation rate. The case *M* = 1 shows, nevertheless, a higher frequency of convergence than expected: under linear ranking, this is the only point in the parameter space we analysed where convergence occurs reliably in so many generations. We hypothesise that, when *M* = 1, slots are statistically allocated in the manner that more closely resembles the expected value than other low *M* values, eliminating some drift. However, further studies and a better statistical insight are necessary to confirm this hypothesis. The effect of *M* can be interpreted as inversely correlated with the variance between an individual’s observed and expected growth curves between generations, according to the GA literature (23).
- A starting population with high frequency (*a*) of a high-quality but unstable strategy (in this case, the putative ancestor strategy; this strategy has a mean pairwise payoff of 1.102, much higher than the average, but only achieves the highest possible payoff against six other strategies) does not slow down convergence to the ES in this size of fitness landscape.

## 4. Discussion

We can think of three measures of the impact of an evolutionary factor on the evolutionary trajectory. We can define, assuming all other conditions fixed: the *speed of convergence* to the global optimum, that is, the average time with which a population converges to it; the *frequency of convergence*, which is the frequency with which populations reach the global optimum, given infinite evolutionary time; and the *accuracy of convergence*, that is, the amount of time, on average, that a population spends with one or more solutions present at very high frequency (*e*.*g*. >80%) in its pool, independently from whether it will ultimately converge to the global optimum or not. Our results show that different assumptions made on the selection model affect the convergence speed. We have also endeavoured to find common patterns in the effect of components of the evolutionary process other than the selection model (*e*.*g*. population size, mutation rate).

We want to stress the limitations of generalising other details of our results to other fitness landscapes. Some fitness landscape features are independent from simplifications that are made at the simulation building stage when studying a biological system: the small size of this landscape and the relatively small difference in fitness between many of its strategies (the ‘flatness’ of the landscape). They nevertheless affect the trajectory of the population over time. Other features we have chosen not to include for simplicity or inapplicability to this game theory landscape: sexual reproduction and hybrid fitness are two of them. They are likely to be major factors influencing the evolutionary trajectory. We believe it to be an interesting future direction to investigate how much of these findings holds true in a larger and differently contoured fitness landscape and under a more complicated relationship between phenotype and genotype. Among the characteristics we suggest testing there are also are the assumption of a continuous landscape and varying mutation sizes.

How important is exploration in reaching the global optimum? Our results show that, the larger the gap in reproductive advantage between the best quality individual and the second best (that is, the greater the exploitation), the faster the convergence, even in a relatively flat landscape. This performance is likely linked to landscape size, relatively to mutation rate and size. This fitness landscape is a high connectivity scenario, because of its small size, meaning that all areas are relatively close together. Therefore, we expect the landscape to have been thoroughly explored by mutation within the 500 generations of our model. High connectivity also means that exploitation is enough to reach the global optimum. In larger landscapes, exploration might play a more important role on the evolutionary endpoint reached, including whether the population can find the global optimum.

The ability of exploitation to find the global optimum is also limited to cases where the optimum is an evolutionary attractor: in the fitness landscape used here, the location with the highest average fitness is also stable (ES), but there are likely to be biological scenarios in which highest average fitness leads to instability and exploration might be essential to reach a stable outcome.

Our investigation focuses on the fitness optimum. However, natural populations are likely to stay in area of high fitness close to but not exactly matching the optimum, due to other constraints including the trade-off between different traits. New analyses should investigate differences between methods in converging to these extended “optimality zones”.

Finally, our results also show that the accuracy of trait quality estimation affects the amount of drift observed. Accordingly, the details of our results are likely to be sensitive to the number of rounds we used in trait evaluation We have provided, in this article, an overview of the methodology of GAs. This modelling framework offers a useful tool to evolutionary biologists interested in the effect of evolutionary process components on a population’s evolutionary trajectory. The advantage of this modelling architecture is that it is already set up to mimic the evolutionary process, thus limiting the degrees of freedom compared to *ad hoc* modifications of a more universal tool like agent-based modelling. This enhances comparability and replicability. GAs, on an equal level to agent-based models, have the flexibility needed for modelling the effect of more complex dynamics, such as the relationship between phenotype and genotype, on the evolutionary outcome. For example, as mentioned, a key feature of our simplistic case study is the absence of phenotypic plasticity. This can easily be integrated into the simulation.

## 5. Conclusion

We have highlighted here the use of genetic algorithms as a simulation framework for studying the effect of all components of the evolutionary process, by presenting a case study of five selection models and their effect on the evolutionary trajectory of a population. Although we emphasise that this outcome cannot be generalised to all possible correlated fitness landscapes, the results outlined here are nevertheless interesting: they highlight substantial differences in the way different assumptions about the evolutionary process influence the expected evolutionary trajectory. We hope that this modelling framework will be used to integrate both known and predicted characteristics of fitness landscapes in the study of evolutionary trajectories. The resulting findings will be useful both to evolutionary theorists and ultimately, if used to generate more accurate models of trajectories for existing populations, to empirical scientists.

## Funding

This work was funded by the John Templeton Foundation as part of the research collaboration grant “Putting the extended evolutionary synthesis to the test” (grant no. 60501) and by the University of St Andrews. Our thanks go to Dr Barbara Trubenova for the keen feedback that has helped reshape this manuscript into a readable form and to Petri Rautiala for essential discussions on the mathematical interpretation of the SPS game landscape.

## Declaration of interests

None.

## Supplementary Information captions

**S1. Derivation of z**

**Supporting File 1. Matrix of payoff resulting from pairwise interaction between all possible strategy pairs.**

## Notes

### Competing Interest Statement

The authors have declared no competing interest.

