## Supplementary Information for "Genetic Algorithms as a method to study adaptive walks in biological landscapes"

### Supporting Information

#### SI.1. Derivation of $z$

Mitchell (1) calculates through integration that

$$Max + min = 2$$

and hence

$$\max(Max) = 2,$$

when the number of reproductive slots *M* coincides with the ranking range *N.* We can generalise this result to any *M*:

$$min + Max = \frac{2M}{N}$$

and, for $Max > min$,

$$Max > \frac{M}{N}.$$

*Max* can be set to a different value from its maximum, *2M/N*, through a parameter *z* similar to *v* in proportional selection, so that $Max = \frac{2zM}{N}$.
